# Pre-determined macromolecular leakage sites with dynamic properties establish endothelial intra-vessel heterogeneity

**DOI:** 10.1101/2020.09.21.305755

**Authors:** M Richards, L Venkatraman, L Claesson-Welsh

**Author notes:** Further information and requests for resources and reagents should be directed to and will be fulfilled by the lead contact, Mark Richards.

## Abstract

Endothelial cells display heterogeneous properties based on location and function. How this heterogeneity influences endothelial barrier stability both between and within vessel subtypes is unexplored. We find here that endothelial cells exhibit heterogeneous barrier properties on inter-organ and intra-vessel levels. Using intravital microscopy and sequential stimulation of the ear dermis with vascular endothelial growth factor-A and/or histamine, we observe distinct, reappearing sites, common for both agonists, where leakage preferentially takes place. Further, through repetitive stimulation of the diaphragm and trachea, we find inter-organ conservation of such leakage sites. Qualitatively, pre-determined sites display distinct leakage properties and enhanced barrier breakdown compared to less susceptible regions. Mechanistically, pre-determined sites exhibit lower laminin α5 deposition, which correlates with reduced junctional vascular endothelial (VE)-Cadherin. These data thus highlight functional intra-vessel heterogeneity that defines pre-determined sites which display distinct leakage properties and which may have great impact on pathological vascular leakage and drug delivery.

## Introduction

The vasculature is restricted from free exchange with surrounding tissues by the vascular barrier composed of endothelial cell (EC) junctions and their surrounding mural cells and basement membrane (BM). The strictness of this barrier can differ between organs with the liver and kidney vasculature possessing endothelial sinusoids and fenestrations conferring elevated permeability (1,2). In organs such as the lungs, muscle and skin, endothelial adherens and tight junctions may widely differ between vessel subtypes in their composition and barrier properties, influencing barrier functionality (3–9) An extreme example is the brain, which is strictly protected from exchange by specialized endothelial tight junctions (10–13).

In response to growth factors such as vascular endothelial growth factor A (VEFGA) and inflammatory cytokines such as histamine, the vascular barrier may be transiently or chronically weakened (14–19). Termed vascular hyperpermeability, this process results in increased leakage of fluid and molecules, as well as increased passage of cells, from the blood. Vascular hyperpermeabilty takes place in solid tumours, under ischemic conditions such as stroke and retinopathies and during inflammation (20–23). In such conditions, uncontrolled and chronic vascular hyperpermeability can lead to edema, tissue ischemia and irreversible tissue damage; it can also hamper standard therapeutic options due to increased interstitial pressure (24). The mechanisms by which vascular hyperpermeability is established is still under much research, however the generally accepted model visualises a paracellular pathway, with the breakdown of adherens junctions and formation of intercellular gaps. The underlying mechanisms involve actin cytoskeleton remodelling and Src family kinase-induced tyrosine phosphorylation of the adherens junction component vascular endothelial cadherin (VE-cadherin), triggering its internalisation (25–29). Organotypic features of vascular hyperpermeability however are far from resolved and have not been studied in a comparative manner. For example, whether different organs respond differently to specific permeability agonists is unclear. Vascular hyperpermeabiltiy also differs within the hierarchy of the vascular system. Over 50 years ago it was proposed that vascular leakage and other such inflammatory functions take place specifically in post-capillary venules (30, 31). Indeed, repetitive stimulation with bradykinin results in reproducible leakage from post-capillary venules (32). However, using the mouse ear dermis as a model, where vessel subtype distinction was made based on hierarchy, blood flow and vessel diameter, we observed through live imaging and intradermal delivery of VEGFA that capillaries were also sensitive to stimulation and underwent vascular leakage, albeit to a lower extent than post-capillary venules (3). Further investigation revealed that within the ear vasculature there exists a gradient of Claudin-5 (Cldn5) expression to which hyperpermeability inversely correlates, from high Cldn5 expression in arterioles to low in venules, whilst intermediate capillaries possess a more variable expression pattern. Furthermore, intravital study of vascular hyperpermeability revealed that bursts of leakage occur at distinct points, whilst the majority of susceptible vessels remain unaffected. Such defined points of disruption likely represent sites in the vessel wall that are more sensitive to stimulation or possess qualities rendering them more susceptible to junctional breakdown.

These observations suggest an endothelial heterogeneity that extends beyond differences within the vascular hierarchy or between organs, to also include inter-EC and even subcellular heterogeneity within the same vessel. Given the advantages afforded by live imaging of vascular leakage we sought to investigate whether such focal intra-vessel heterogeneity does indeed exist and whether phenotypic differences are present within the same vessel. Through repetitive stimulation of the mouse ear dermis with VEGFA and/or histamine we observe a high proportion of leakage takes place recurrently from distinct sites, which are common to both ligands. Such pre-determined sites of leakage are also found to exist in both the trachea and diaphragm. In depth analysis of their dynamic properties demonstrates that pre-determined sites exhibit enhanced barrier disruption and leakage rate compared to less susceptible sites of leakage. Such enhanced barrier disruption correlates with reduced deposition of laminin α5 and lower junctional VE-Cadherin. Thus, we show that within the microvasculature there exists pre-determined sites of leakage and demonstrate a heterogeneity of the vasculature that goes beyond organotypicity and vessel subtype variability to reveal phenotypic differences between adjacent ECs of the same vessel.

## Results

### VEGFA-induced leakage takes place from pre-determined sites

We recently reported the development of a technique that allows the live visualisation of leakage from the mouse ear microvasculature in response to the abluminal delivery of factors such as VEGFA in an atraumatic manner (Figure 1A) (3). Using this technique, it became evident that leakage in response to VEGFA takes place from geographically distinct positions, signifying the restricted localisation of barrier disruption to precise sites (approximately 1-2 μm in diameter), as opposed to widespread loosening of the endothelium or breakdown of an ECs entire junctional perimeter. Further exploration showed that these sites may often be distant from one another and independent of the distance from the ligand injection site (Figure 1B and Movie 1). Thus, even though a vessel may have the capacity to, or be expected to leak (e.g a post-capillary venule), it is not necessarily the case that it will leak, or that leakage will occur uniformly along its entire length. Consequently, ECs and subcellular sites within vessels may display very different susceptibilities to stimulation.

**Figure 1.**
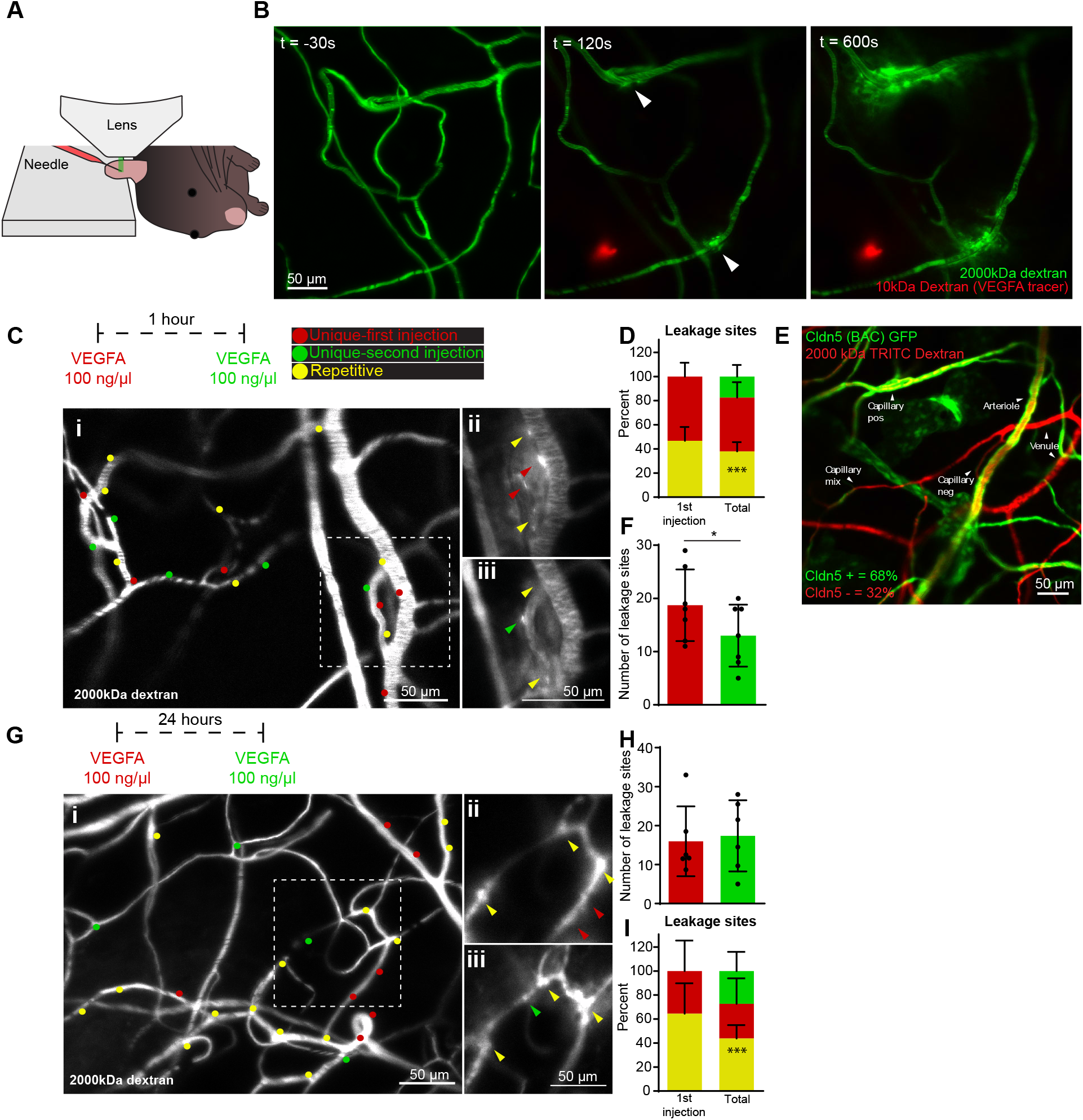
Pre-determined sites of leakage in response to VEGFA. **A.** Schematic representation of intravital imaging of the mouse ear dermis coupled with intradermal microinjection. **B.** Stills showing vascular leakage in response to VEGFA injection. Arrowheads mark sites of vascular leakage. **C.** Representative image of leakage sites in response to repetitive VEGFA injection with a 1 hour interval. Red and green dots show unique sites following first and second injection of VEGFA respectively, yellow dots show repetitive sites. Boxed region is shown as magnified inserts to the right and shows leakage sites (coloured arrowheads) following the first (i) and second injection (ii) (See Movie 2). **D.** Quantification of leakage sites; repetitive (yellow), unique 1st (red), unique 2nd (green), as a percentage of those seen following the first injection (left) or of both injections (total; right) of VEGFA, with a 1 hour interval. n = 7 mice, ≥ 2 acquisitions/mouse, p < 0.001. **E.** Representative image of Cldn5 expression in the ear microvasculature as shown using a reporter mouse expressing GFP under the Cldn5 promoter (Cldn5 (BAC) GFP). Cldn5 positive cells are shown in green, Cldn5 negative vessels in red. n = 5 mice. **F.** Number of leakage sites following the first and second injection of VEGFA, with a 1 hour interval. n = 7 mice, ≥ 2 acquisitions/mouse, p = 0.0287. **G.** Representative image of leakage sites in response to repetitive VEGFA injection with a 24 hour interval. Red and green dots show unique sites following first and second injections respectively, yellow dots show repetitive sites. Boxed region is shown as magnified inserts to the right and shows leakage sites (coloured arrowheads) following the first (i) and second injection (ii). **H.** Number of leakage sites following the first and second injections of VEGFA with a 24 hour interval. n = 6 mice, ≥ 2 acquisitions/mouse. **I.** Quantification of leakage sites; repetitive (yellow), unique 1st (red), unique 2nd (green), as a percentage of those seen following the first injection (left) or of both injections (total) of VEGFA, with a 24 hour interval. n = 6 mice, ≥ 2 acquisitions/mouse, p < 0.001. Error bars represent mean and SD. Statistical significance was determined using a two-tailed paired or unpaired Students’ t-test where appropriate. See also Figure S1 and S3.

To ascertain whether certain sites on blood vessels have a higher sensitivity to stimulation, or are pre-determined to leak, VEGFA injection of the same microvascular region was carried out on two consecutive occasions using an interval of 1 hour. Sites of leakage were visualized by extravasation of circulating 2000 kDa fluorescent dextran and were defined as distinct leakage sites when occurring at least 5 μm apart along the longitudinal axis of the vessel, to allow for clear separation of individual bursts of leakage. Analysing the reproducibility of leakage from vessel segments (between bifurcations) showed a high reproducibility of leakage, up to 75%, within a typical vessel stretch of approximately 200 μm (Figure S1), similar to previously published results (32). Interestingly, higher resolution analysis demonstrated that 47% of precise subcellular sites of leakage during the first injection of VEGFA also leaked following the second (i.e. leakage was seen within 5 μm of a previously observed site) (Figure 1C-D and Movie 2). In general these sites occurred once every 100 μm of a permissible vessel (Figure S2). Importantly, these repetitive sites shared similar distance profiles, relative to the site of VEGFA injection, as unique sites that only leaked following one injection (first or second), thus ruling out repetitive sites occurring due to higher VEGFA presentation at the endothelial surface (Figure S3A).

Compared to the total number of leakage sites seen following both injections, repetitive leakage sites represented 38% (Figure 1D). To assess whether this might constitute a significant difference compared to random leakage site-selection, we employed a previously characterised mouse model where VEGFA-resistant vessel segments are marked by green fluorescent protein (GFP) reporting the expression of Cldn5 (Cldn5 (BAC) GFP) (3). Of the ear microvasculature, 32% was found to be Cldn5 negative and therefore permissive to leakage (Figure 1E). Given the average length of blood vessels within imaged fields (5,800 μm) and assuming 32% of this length was Cldn5 negative (1,856 μm), random leakage site-selection would produce repetitive leakage sites at a rate of 0.4% at any given point (with a tolerance of 5 μm producing 371 potential sites and assuming binomial distribution). Compared to the actual rate of 38%, observed repetitive leakage sites are thus significantly unlikely to occur by chance alone.

Stimulation of ECs with VEGFA commonly results in cell desensitisation (33). Here, comparison of leakage site number following first and second injections showed a significant decrease of 31% subsequent to second stimulations (Figure 1F). Thus EC desensitisation may affect the leakage site proportions than would otherwise be observed.

To assess whether this desensitisation affects the proportion of unique and repetitive leakage sites, repeated VEGFA injections were carried out with an interval of 24 hours, to allow fuller recovery of ECs. As expected, a longer recovery period resulted in an increase in the proportion of second versus first injection leakage sites, with 65% of sites that leaked following the first injection also leaking following the second (Figure 1G-H). Further, with a 24 hour interval in-between injections, repetitive leakage sites were 44% of the total, i.e. a significant increase compared to random leakage site-selection (Figure 1G-I).

Thus, this data shows that within the microvasculature of the ear dermis there exists sites that exhibit a higher susceptibility to leakage in response to VEGFA and highlight an intra-vessel functional heterogeneity.

### Pre-determined sites of leakage are conserved between permeability agonists

Vascular leakage may be induced by a number of factors, with cell surface receptor tyrosine kinases or G-protein coupled receptors directly or indirectly regulating intracellular signalling leading to junctional disruption (14). To investigate whether the existence of pre-determined sites of leakage is specific to VEGFA, or a more general feature of EC biology, the inflammatory cytokine histamine was similarly investigated for its pattern of leakage induction.

Similar to VEGFA, abluminal histamine stimulation resulted in junctional breakdown and bursts of leakage at distinct sites. Repeated histamine injection in the ear dermis with a 1 hour interval showed that 30% of the sites that leaked following the first injection also leaked following the second (Figure 2A-B). These repetitive sites represented 20% of the total sites seen, which is a lower proportion than observed for VEGFA but still a significant increase in comparison to random leakage site-selection. Further, distance profiles from the site of injection of repetitive and unique sites were found to be the same, ruling out any effects from higher histamine concentrations nearer the site of injection (Figure S3B).

**Figure 2.**
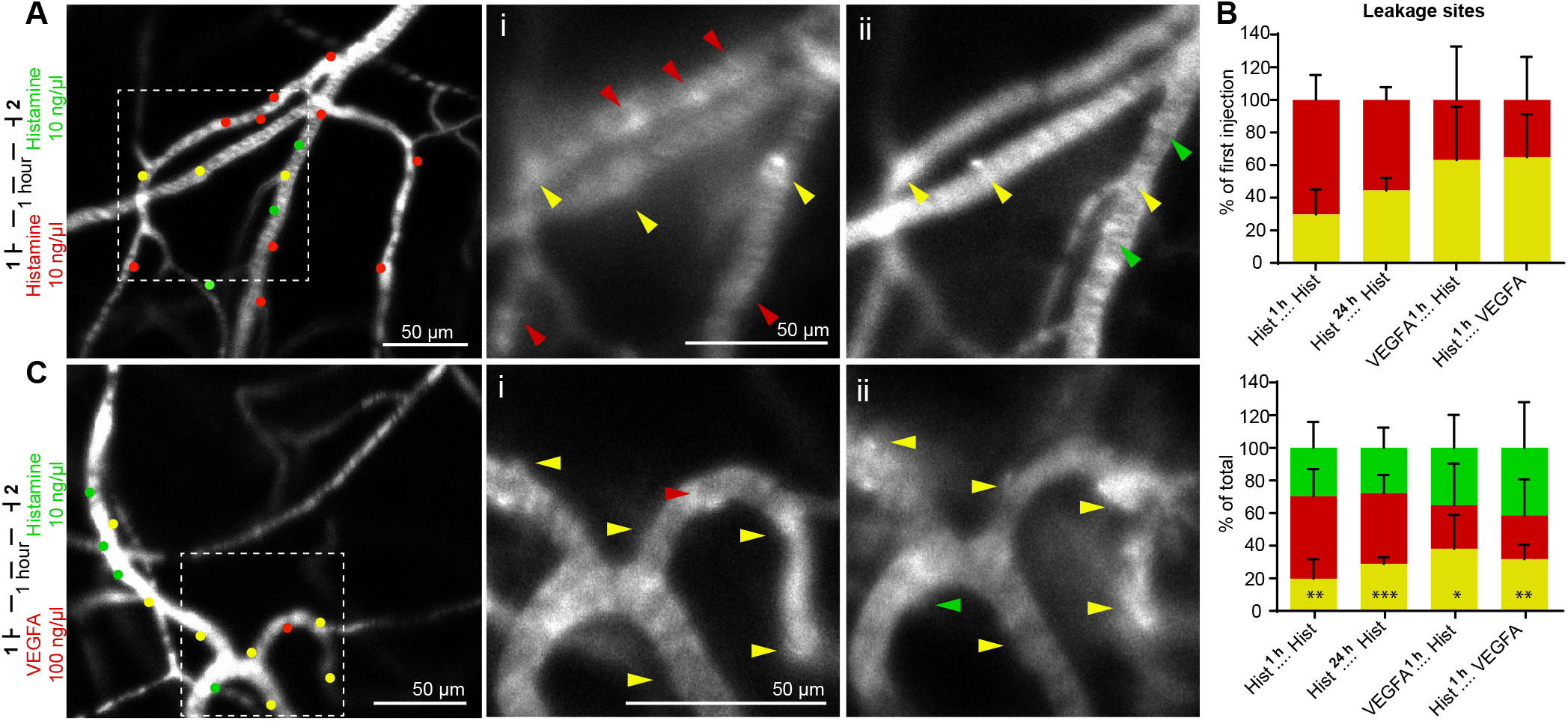
VEGFA and histamine induce leakage from the same pre-determined sites. **A.** Representative image of leakage sites in response to repeated histamine injections with a 1 hour interval. Red and green dots show unique sites following first and second injection respectively, yellow dots show repetitive sites. Boxed region is magnified to the right and shows leakage sites (coloured arrowheads) following the first (i) and second injection (ii). **B.** Quantification of leakage sites; repetitive (yellow), unique 1st (red), unique 2nd (green), as a percentage of those seen following the first injection (top) or of both injections (bottom) of histamine with a 1 or 24 hour interval (p = 0.0029 and p < 0.001 respectively), VEGFA followed by histamine (p = 0.0362) or histamine followed by VEGFA (p = 0.0015) (1 hour intervals). n ≥ 4, ≥ 2 acquisitions/mouse. **C.** Representative image of leakage sites in response to VEGFA followed by histamine with a 1 hour interval. Red and green dots show unique sites following first and second injections respectively, yellow dots show repetitive sites. Boxed region is magnified to the right and shows leakage sites (coloured arrowheads) following the first (i) and second injection (ii). Error bars represent mean and SD. Statistical significance was determined using a two-tailed unpaired Students’ t-test. See also Figure S2 and S3.

Decreased repetitive sites compared to VEGFA could be due to a stronger desensitisation following histamine injection, as evidenced by 42% fewer sites occurring following the second injection (Figure S3E); compared to a 31% reduction with VEGFA (Figure 1F). To explore this, repeated histamine injections were also carried out with an interval of 24 hours. Similar to VEGFA, a longer interval period resulted in increased repetitive sites, with 45% of those sites seen following the first injection also leaking following the second (Figure 2B-C). Meanwhile, repetitive leakage sites represented 29% of the total seen, significantly more than if selection were random (Figure 2B). Therefore, a subset of leakage sites in response to histamine are also pre-determined, although perhaps to a lower degree than VEGFA. However, even with an interval of 24 hours, a second histamine injection produced 31% fewer leakage sites than the first (Figure S3E), whilst an interval of 24 hours restored endothelial sensitivity to VEGFA (Figure 1I). Thus, the decreased incidence of repetitive leakage sites in response to histamine may be due to a stronger and more sustained refractory period.

The existence of pre-determined sites of leakage in response to both VEGFA and histamine led us to investigate whether these sites may in fact be the same for both ligands, and whether certain vessel sites in general exhibit a higher susceptibility to leakage. To achieve this, we visualised vascular leakage in response to VEGFA or histamine, followed by histamine or VEGFA respectively. Using an interval of 1 hour, it was found that VEGFA and histamine induced leakage from the same sites, to a high degree. Stimulation with VEGFA followed by histamine, or histamine followed by VEGFA, resulted in repetitive leakage sites representing 63% and 65% of first injection leakage sites respectively and 35-40% of the total seen (Figure 2B-C). Any effects resulting from differences in ligand potency were negated at the concentrations used, as evidenced by the similar number of leakage sites induced by both ligands and the similar distance profiles of sites induced by VEGFA or histamine (Figure S2 and S3C-D). Similarly, there was no apparent effect of endothelial desensitisation, even given the short 1 hour interval period (Figure S3E).

Thus, these data demonstrate that beyond leakage being specific to post-capillary venules, as previously described, or specific to Cldn5-negative vessels as we have recently shown (3), a significant proportion of leakage takes place from distinct subcellular sites on the endothelium that are pre-determined, where sensitivity to stimulation is higher and repetitive leakage takes place more readily.

### Distinct properties of repetitive and unique sites

Given our observation that a subset of sites within the ear microvasculature have a pre-determined capacity to leak, we investigated whether these repetitive sites differed in leakage properties compared to unique sites. Through the real-time visualisation of vascular leakage we can quantify factors such as the lag period (how long it takes ECs to respond to stimulation), and also the degree of leakage (the rate of extravascular dextran appearance at sites of leakage), which tells us the kinetics by which dextran appears outside the vessel and gives us an indication of the degree of localised endothelial barrier disruption (Figure 3A).

**Figure 3.**
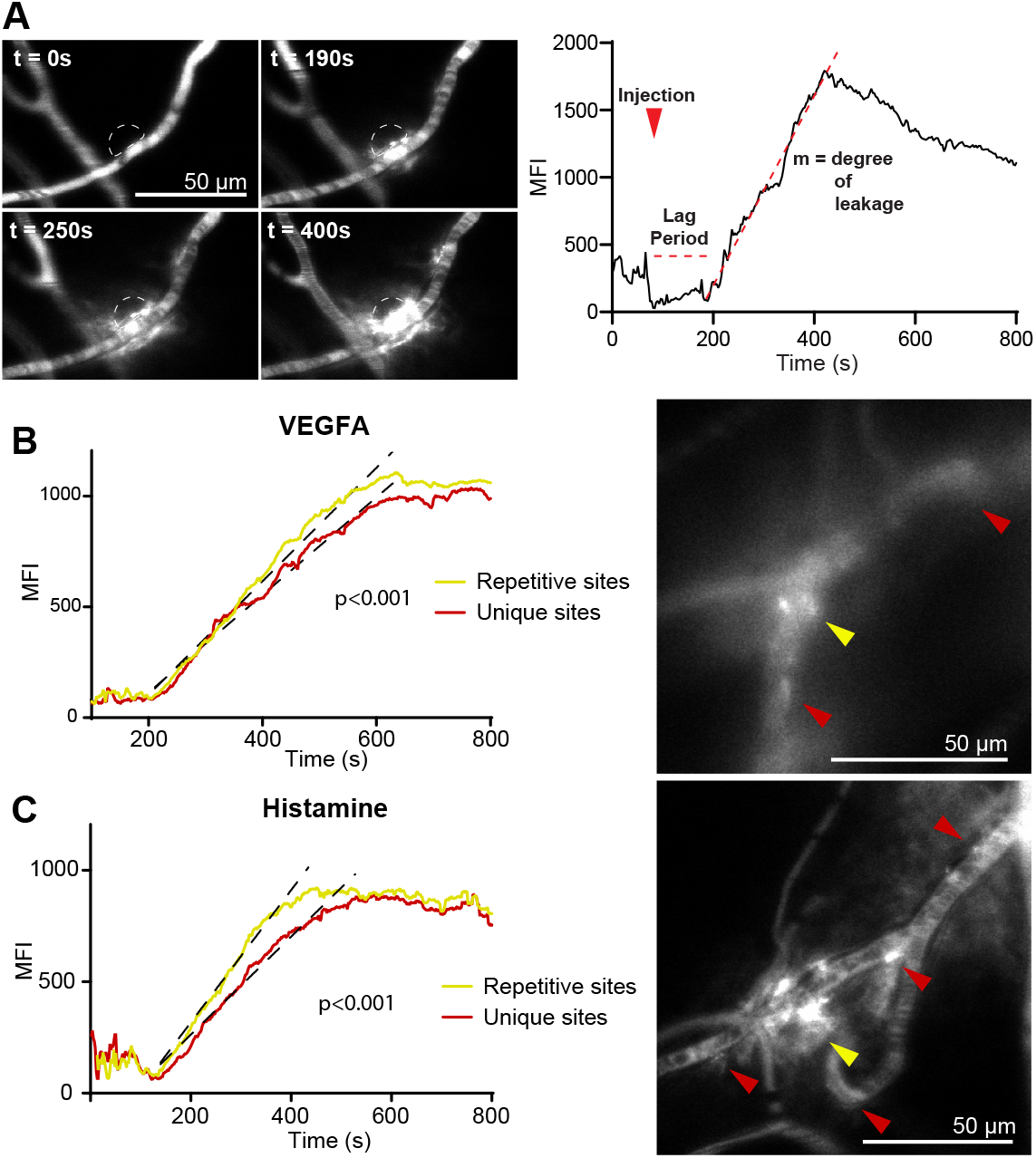
Pre-determined sites of leakage exhibit distinct leakage profiles. **A.** Representation of lag period and degree of leakage analysis. Quantification of extravascular change in fluorescence intensity at site of leakage (dashed region) allows calculation of the time between agonist injection and initiation of leakage (lag period) and the rate of dextran extravasation (gradient, m) (degree of leakage). **B.** Left. Line profiles of the change in fluorescence intensity at sites of leakage in response to VEGFA injection. Lines of best fit for the slopes between leakage initiation and leakage termination are shown with black lines. Right. Representative image of leakage at repetitive and unique sites in response to VEGFA (coloured arrowheads). n = 16, ≥ 2 acquisitions/mouse. **C.** Left. Line profiles of the change in fluorescence intensity at sites of leakage in response to histamine injection. Lines of best fit for the slopes between leakage initiation and leakage termination are shown with black lines. Right. Representative image of leakage at repetitive and unique sites in response to histamine (coloured arrowheads). n = 16, ≥ 2 acquisitions/mouse. MFI, mean fluorescence intensity. Statistical analysis was carried out using linear regression and ANCOVA. See also Figure S4.

Quantification of the lag period demonstrated that repetitive and unique sites generally did not differ in this aspect, with both starting to leak 3-5 minutes after stimulation with VEGFA or histamine (Figure S4A). Of note, unique sites occasionally displayed a significantly higher variability in these dynamics, indicating that signalling at repetitive sites may follow a more consistent pattern than that of unique sites. Analysis of the degree of leakage, comparing unique and repetitive sites following initial VEGFA or histamine injection, revealed that unique sites leak at a significantly lower rate than repetitive sites (Figure 3B-C and Movies 3-4). Similarly, examination of datasets from subsequent second injections showed that this held true for all conditions except VEGFA injections with a 1 hour interval, where leakage occurred at a similar rate for repetitive and unique leakage sites (Figure S4B).

This thus shows that at unique sites, the endothelium undergoes less disruption resulting in a slower extravascular release of dextran, whereas at repetitive sites, a stronger response leads to enhanced barrier disruption, larger intercellular gap formation and greater macromolecular extravasation.

### Low laminin α5 deposition marks pre-determined sites of leakage

To explore vessel properties potentially defining sites of pre-determined leakage we developed methodology to allow accurate localisation of leakage sites after fixation and immunofluorescent staining. To this end fluorescent microspheres with distinct excitation and emission spectra were combined with consecutive tail-vein histamine stimulations (Figure 4A). Similar to the live imaging studies, tail-vein histamine stimulation (12.5 mg/kg) produced a sustained refractory period (Movies 5-6). Thus a 24 hour interval was necessary to allow re-sensitisation and leakage to occur. Given this long interval period 200 nm microspheres were chosen based on their ability to pass through interendothelial gaps but likeliness to remain largely *in situ*.

**Figure 4.**
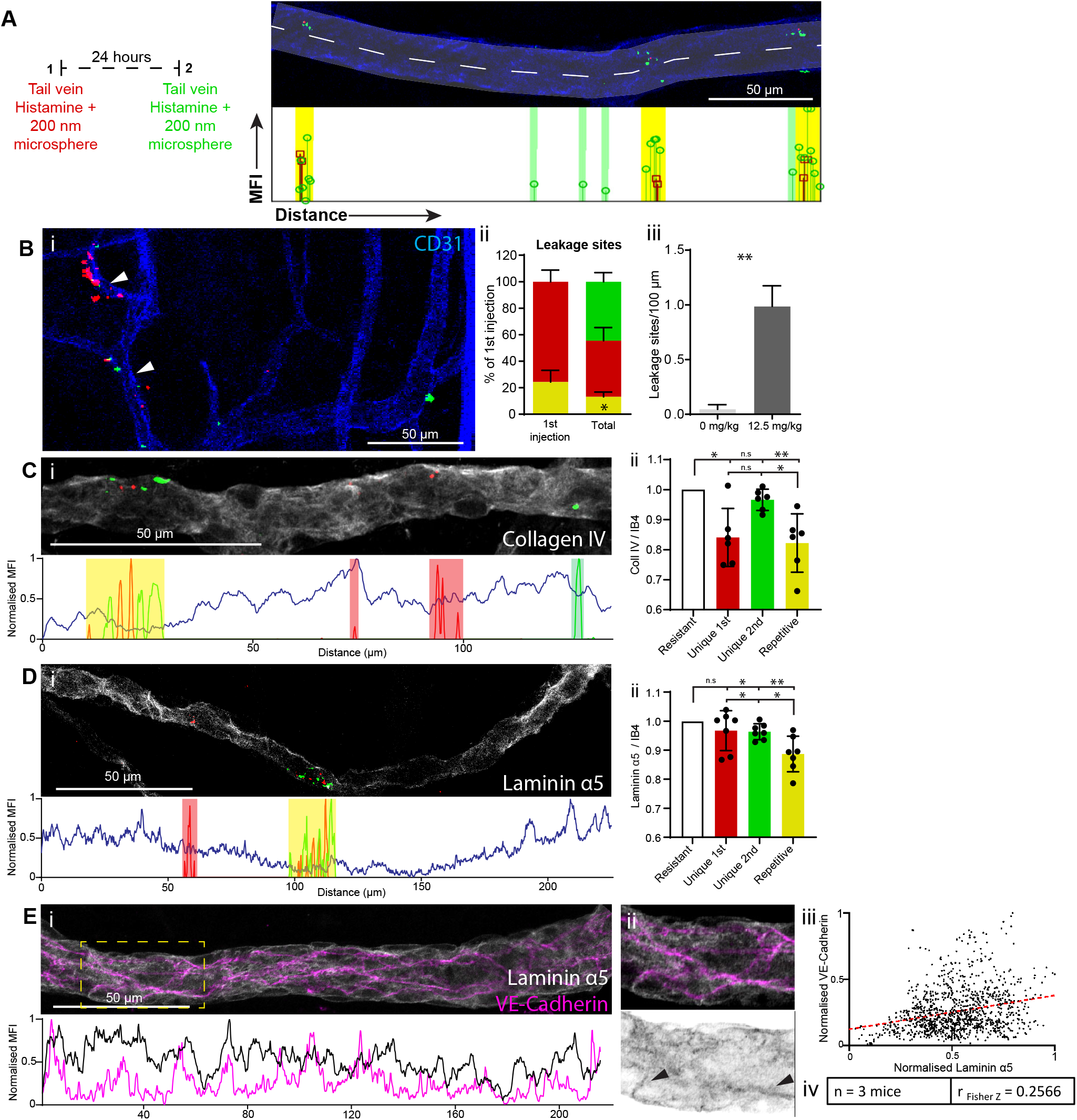
Low laminin α5 deposition marks pre-determined sites of leakage. **A.** Schematic showing experimental design (left) and analysis (right). Extravasated microspheres are quantified through fluorescence analysis along the long axis of vessels (dashed line) and leakage sites classified as unique or repetitive according to the location and overlap of different microspheres. Lollipops in red and green represent extravasated microspheres from the first and second injection respectively. Lollipop length represents mean fluorescence intensity (MFI) of microspheres at their site of leakage. Green and yellow bars highlight those sites classified as unique (for second injection) and repetitive respectively. **B.** i. Representative image of microsphere leakage in the mouse ear in response to circulating histamine. Arrowheads show sites of repetitive leakage. ii, Bar chart showing quantification of leakage sites; repetitive (yellow), unique 1st (red), unique 2nd (green), as a percentage of those seen following the first injection (left) or of both injections (right). p = 0.0406, comparing the proportion of repetitive leakage sites if site-selection were random. iii, Bar chart showing leakage sites per vessel length with and without histamine stimulation, p = 0.0011, average of 43 mm of vessels analysed / mouse. **C.** i. Representative image (Above) and its long-axis quantification (Below) of collagen IV correlation with microsphere leakage. ii. Bar chart showing comparison of collagen IV / IB4 intensity within vessel segments at leakage sites. Intensities are normalised to collagen IV / IB4 at sites within segments where no leakage was seen (Resistant). Compared to Resistant: Unique 1st; p = 0.0099, Unique 2nd; p = 0.0648, Repetitive; p = 0.0065. Compared to Repetitive: Unique 1st; p = 0.7311, Unique 2nd; p = 0.0109. n = 6, ≥ 3 fields of view / mouse, ≥ 6 vessel segments / field of view. **D.** i. Representative image (Above) and its long-axis quantification (Below) of laminin α5 correlation with microsphere leakage. ii. Bar chart showing comparison of laminin α5 / IB4 intensity within vessel segments at leakage sites. Intensities are normalised to laminin α5 / IB4 at sites within segments where no leakage was seen (Resistant). Compared to Resistant: Unique 1st; p = 0.253, Unique 2nd; p = 0.0128, Repetitive; p = 0.0028. Compared to Repetitive: Unique 1st; p = 0.0112, Unique 2nd; p = 0.0486. n = 7, ≥ 3 fields of view / mouse, ≥ 6 vessel segments / field of view. **E.** i. Representative image (Above) and its long-axis quantification (Below) of laminin α5 and VE-Cadherin in a post-capillary venule. VE-Cadherin quantification is normalised to the number of junctional elements within the vessel cross-section. ii. Magnification of insert in i. Arrowheads highlight regions of linear, junctional laminin α5. iii. Representative scatter dot-plot and line of best fit obtained through quantification of VE-Cadherin and laminin α5 along the long-axis of the vessel in i. iv. Table summarising data from n = 3 mice, with ≥ 6 fields of view / mouse and ≥ 3 vessels / field of view. Error bars represent mean and SD. Statistical significance was determined using a two-tailed unpaired or paired Students’ t-test. Correlation was calculated using Spearman’s rank-order test and Fisher’s Z transformation. See also Figure S5.

Following repeated histamine and microsphere administration, mice were perfused and ear vasculature imaged. Leakage site categorisation as unique (red or green, following first and second injection, respectively) or repetitive (yellow) was achieved following fluorescence analysis along the vessel longitudinal axis (Figure 4A and Methods). These analyses revealed that microsphere leakage from ear dermal vasculature following repetitive histamine stimulation often took place from the same, pre-determined sites (Figure 4B), similar to our findings when assessing leakage of circulating 2000 kDa dextran in response to intradermal injection of histamine (Figure 2). Quantification showed that 25% of those sites that leaked following the first injection and 14% of the total number of leakage sites seen were classified as repetitive, a significant increase over what would be expected from random leakage site-selection (Figure 4Bii). Importantly, the similar number of observed red (first injection) and green (second injection) leakage sites demonstrated that microsphere movement from sites of leakage over the 24 hour interval was minimal. Furthermore, the specificity of identified leakage sites as being induced by histamine, as opposed to microspheres passively passing the endothelial barrier, was shown through comparison with unstimulated tissue (Figure 4Biii). Thus, consecutive administration of fluorescent microspheres and histamine allows accurate localisation of unique and repetitive leakage sites.

An important regulator of EC properties is the underlying BM. During the inflammatory process leukocyte transmigration correlates with low expressing regions of BM components including collagen IV and laminin α5 (34, 35). Such patchy expression can be seen in post-capillary venules of the ear dermis, whereas arterioles exhibit a uniform high expression (Figure S5). In comparison capillaries exhibit a low but uniform expression.

Low expressing regions are believed to offer the path of least resistance for cell passage across the BM. Levels of collagen IV and laminin α5 however have also been found to regulate endothelial junction properties through modulation of VE-Cadherin and Claudin-5 expression and localisation (36, 37). Thus, these BM components were investigated for their expression levels at sites of repetitive and unique leakage, as well as regions where no leakage was observed (resistant). Immunofluorescent staining for collagen IV showed that whilst its expression was lower at sites of repetitive leakage compared to resistant sites, this was also true for unique sites following the first stimulation (Figure 4C). Unique sites from the second stimulation however showed no difference compared to resistant vessel sites. Laminin α5 meanwhile showed a specific and significant reduction in expression at sites of repetitive leakage compared to resistant and unique sites (Figure 4D). Thus, BM laminin α5 levels inversely correlate with endothelial response to stimulation, with enhanced barrier disruption occurring at laminin α5 low sites.

VE-Cadherin junctions are disrupted in laminin α5 knock-out mice whilst culture of bEnd.3 cells on laminin α5 enhances junctional VE-Cadherin localisation (36). We thus wished to explore whether the low laminin α5 expression at pre-determined sites of leakage exhibited particular VE-Cadherin morphology. As junctions are likely to be affected by stimulation and leakage, analysis at sites of leakage after agonist injection, was not suitable. We thus directly analysed and correlated laminin α5 deposition with underlying VE-Cadherin junctions. Analysing post-capillary venules (diameter 10-30 μm), which consistently show susceptibility to leakage, we identified a weak positive correlation between junctional VE-Cadherin concentration and laminin α5 deposition (r_Fisher’s Z_ = 0.2566) (Figure 4E). Moreover, dense, fibrillar regions of laminin α5 deposition were often seen to overlap with VE-Cadherin junctions (Figure 4Eii). These data thus highlight a correlation between VE-Cadherin localisation and laminin α5 deposition within the BM. Such regions, expressing low levels of laminin α5 and VE-Cadherin, may constitute sites of pre-determined leakage with a higher susceptibility to undergo barrier breakdown in response to stimulation.

### Pre-determined sites of leakage are conserved between vascular beds

Given our findings that dermal blood vessels have pre-determined sites of leakage we wanted to investigate whether other vascular beds possess similar qualities.

Dermal blood vessels are built up of non-fenestrated continuous endothelium, where dogma predicts that induced macromolecular leakage takes place following formation of intercellular gaps (30, 38, 39). We thus wished to analyse organs also possessing continuous endothelium. Given the possibility of relatively simple whole mounting and imaging of the mouse trachea and diaphragm, these organs were chosen for further study. As these organs are difficult to access, atraumatic live imaging is not possible and thus we utilised consecutive tail-vein administration of fluorescent microspheres to localise sites of leakage.

Leakage in the trachea, although still specific to certain vascular segments, was much stronger in response to histamine than that seen in the ear, (Figure S6), possibly reflecting the immune surveillance function of the airway. This strong permeability response precluded the identification of individual sites of leakage. However the use of lower concentrations of histamine (0.5 mg/kg) led to leakage from more distinct sites, representing those with a higher sensitivity to stimulation. Investigation of whether these sites might represent pre-determined sites showed that similar to the ear dermis, repetitive leakage sites represented 27% of those seen following the first stimulation and 21% of the total (Figure 5Ai-ii). A larger number of red (first injection) than green (second injection) leakage sites was observed potentially indicating desensitisation even at 24 hours; but persistently, injection of microspheres alone resulted in negligible leakage also in the trachea (Figure 5Aii-iii).

**Figure 5.**
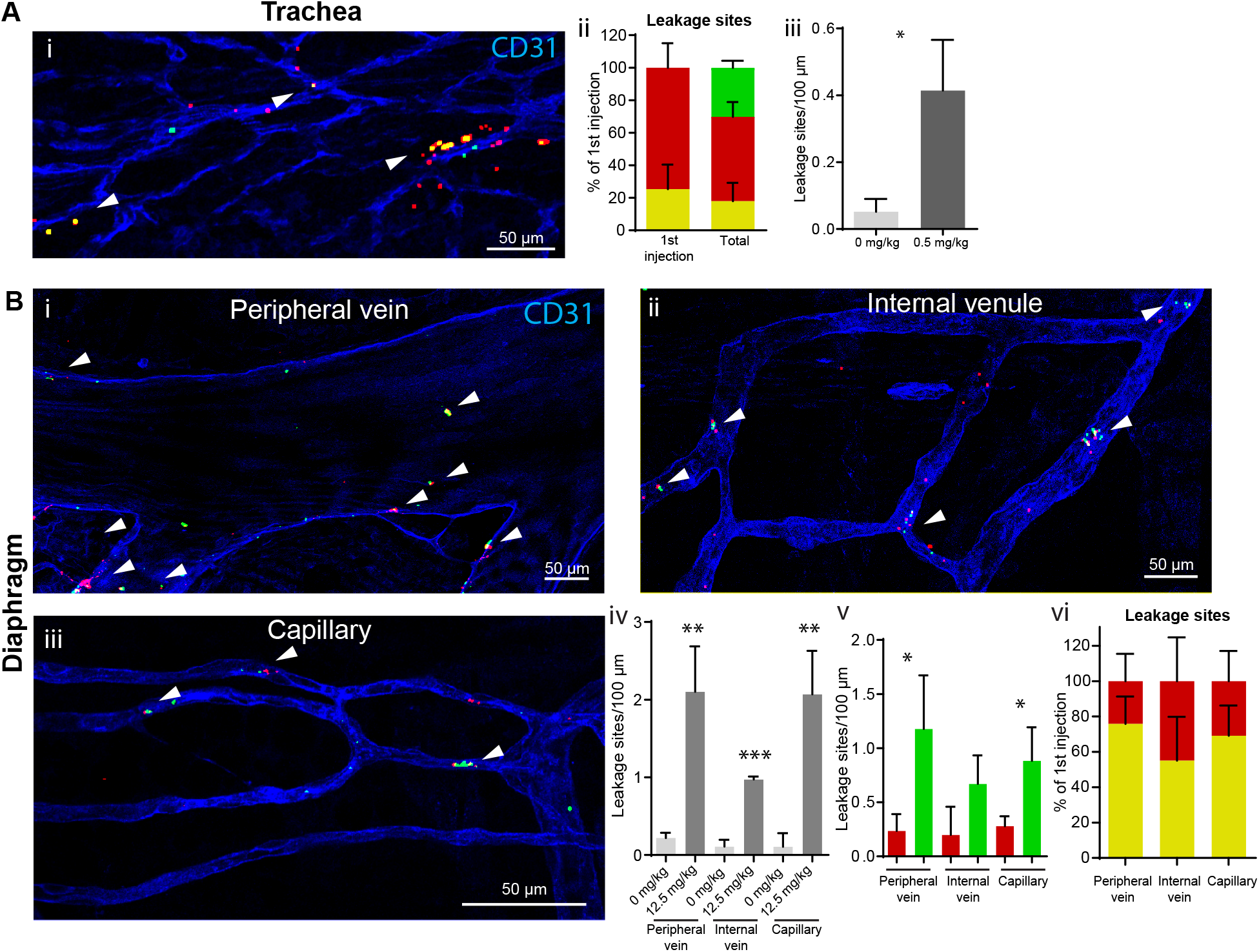
Pre-determined sites of leakage are conserved between vascular beds. **A.** i. Representative image of microsphere leakage in the mouse trachea in response to circulating histamine. Arrowheads show sites of repetitive leakage. ii, Bar chart showing quantification of leakage sites; repetitive (yellow), unique 1st (red), unique 2nd (green), as a percentage of those seen following the first injection (left) or of both injections (right). iii, Bar chart showing leakage sites per vessel length with and without histamine stimulation, p = 0.0159, average of 76 mm of vessels analysed/mouse **B.** i-iii. Representative images of microsphere leakage from the peripheral vein (i), internal venules (ii) and capillaries (iii) of the mouse diaphragm in response to circulating histamine. Arrowheads show sites of repetitive leakage. iv, Bar chart showing leakage sites per vessel length with or without histamine stimulation; (peripheral vein, p = 0.0091; internal vein, p < 0.001; capillary, p = 0.0089). v, Bar chart showing quantification of leakage sites per vessel length in response to first (red) and second (green) histamine injections; (peripheral vein, p = 0.0343; capillary, p = 0.0317). vi, Bar chart showing quantification of leakage sites; repetitive (yellow), unique 1st (red), as a percentage of those seen following the first injection. n = 3, average of 20 mm of vessels analysed/mouse. Error bars represent mean and SD. Statistical significance was determined using a two-tailed unpaired Students’ t-test. See also Figure S6.

The diaphragm vasculature consists of a large calibre peripheral vein lying adjacent to the muscle as well as venules and capillaries within the central tendon. Similar to the ear vasculature, individual leakage sites in the diaphragm were observed using 12.5mg/kg histamine and leakage sites were largely specific to histamine induction (Figure 5Bi-iv). Conversely, unlike the dermis and trachea, a large decrease in the number of red (first injection) compared to green (second injection) microsphere leakage sites was seen, possibly due to organotypic effects resulting in microsphere diffusion during the 24 hour interval (Figure 5Bv). Nevertheless, when compared with those observable sites that leaked following the first injection, repetitive leakage sites represented 75%, 55% and 69% in peripheral veins, internal venules and capillaries respectively (Figure 5Bvi).

These data thus not only show organotypic differences with regard to EC sensitivity and vessel structure, but that intra-vessel heterogeneity also exists in multiple organs and may perhaps also be a general feature of the entire vascular system.

## Discussion

Here we report that distinct subcellular sites of vascular leakage are pre-determined and reappear in response to repeated injection with permeability agonists in the dermis, trachea and diaphragm. Moreover, to a considerable extent, identical sites are shared between VEGFA and histamine. Previous publications have reported the restricted and repetitive vascular leakage from post-capillary venules, which we previously further defined to include pre-venular capillaries and post-capillary venules lacking Cldn5 expression (3, 31, 32). We now show with higher resolution imaging and analysis that within these permissive vessels, inter-endothelial and subcellular heterogeneity gives endothelial junctions that are more sensitive to stimulation than other adjacent, seemingly identical segments of the microvasculature. Additionally, such pre-determined leakage sites undergo more extensive disruption than more resistant, unique leakage sites.

Sites of preferential leukocyte extravasation have previously been associated with decreased levels of basal laminin α5 and collagen IV (34, 35). Here we find that pre-determined sites of macromolecular leakage are marked by a specific decrease in laminin α5 but not collagen IV. Collagen IV is reduced at unique sites that leak only following the first stimulation and at pre-determined sites, suggesting that subsequent to leakage, collagen IV is degraded at these sites. Interestingly, sites that leak only following the second stimulation, showed higher levels of collagen IV, demonstrating specific properties of these different leakage site categories. Laminin α5 meanwhile is specifically reduced at pre-determined, but not unique, sites of leakage and this correlates with decreased local junctional VE-Cadherin concentration. Overall this work highlights an intra-vessel heterogeneous deposition of laminin α5 that impacts localised VE-Cadherin junctional concentration, facilitating barrier breakdown at pre-determined sites (Figure 6).

**Figure 6.**
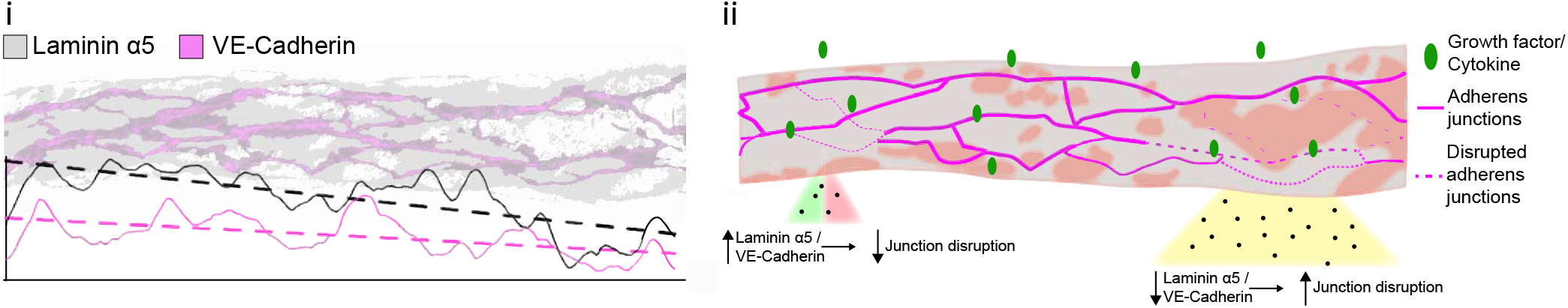
Schematic representation of mechanism. Schematic representation of proposed mechanism underlying pre-determined sites of leakage. i. Laminin α5 regulates junctional localisation of VE-Cadherin with sites of lower Laminin α5 correlating with finer, weaker adherens junctions. Graph below illustrates laminin α5 and junctional VE-Cadherin intensities with lines of best fit. ii. Exposure of vessels to stimulatory factors such as growth factors and cytokines leads to a heterogeneous leakage response. Pre-determined sites of leakage (Right) have lower laminin α5 and thus junctional VE-Cadherin predisposing to enhanced barrier disruption in a repetitive manner. Meanwhile unique or resistant sites (Left) exhibit higher laminin α5 and junctional VE-Cadherin resulting in a diminished or absent leakage response.

The implication of laminin α5 in defining both sites of macromolecular leakage and leukocyte extravasation raises the question of whether these are shared. However, pre-determined sites of leakage often exist in vessels as small as 8 μm in diameter, far smaller than those from which leukocyte extravasation is observed to occur. Further, sites of leukocyte extravasation have been found to have a 60% reduction in the expression of laminin α5 and collagen IV (34); in contrast, predetermined sites of leakage exhibit specific reduction in laminin α5 only and a more modest, although highly significant reduction of 12%, compared to resistant sites. Thus, pre-determined sites of leakage appear unlikely to be amenable to leukocyte passage.

From the data presented here, and by others (36), it is tempting to speculate that levels of junctional VE-Cadherin alone may define sites of preferential leakage and leukocyte extravasation. Notably, reduced levels of VE-Cadherin at subcellular sites drives junctional dynamics through the induction of asymmetrical cytoskeletal dynamics and formation of junction associated intermittent lamellipodia (JAILs) (40). Inhibition of such structures has been associated with loss of barrier integrity in response to thrombin. The formation of such subcellular sites of junctional heterogeneity is in keeping with our observations, where the precise sites of leakage (±5 μm) constitute a fraction of the space occupied by an EC. However, targeting VE-Cadherin *in vivo* through blocking antibodies or its induced deletion leads to enhanced vascular leak in only a subset of organs such as the heart and lungs that are exposed to high mechanical stress (41, 42). VE-Cadherin may also influence endothelial junction stability depending on its phosphorylation status downstream of intracellular kinases such as Src and phosphatases such as VE-PTP (26, 27, 43–48). Whether heterogeneous deposition of BM components might impact VE-Cadherin phosphorylation patterning to generate sites of preferential leakage or leukocyte extravasation is yet to be uncovered.

Beyond direct effects on VE-Cadherin, BM composition and structure may impact junctional dynamics in a number of different ways. For example, BM stiffness correlates with EC contractility, with stiffer substrates allowing exertion of greater RhoA-ROCK based contraction and greater junctional destabilisation (49, 50). Heterogeneous BM structure may thus define localised ‘hot spots’ of leakage, in agreement with the data presented here. Further, through outside-in integrin signalling the BM impacts on the vascular barrier. In particular, the actin cytoskeleton acts as an important link between integrins and endothelial junction stability. Through α- and β-catenin, VE-Cadherin junctions are coupled to the actin cytoskeleton (51, 52). There are also direct links between integrins β1, β3 and β5, the actin cytoskeleton and its effectors during barrier disruption (53–55). Intriguingly, organotypic barrier effects downstream of such integrins has been reported and likely reflects the variability in BM composition between organs (54, 56). It is thus likely that variable BM patterning which we also find here, has wide implications on cytoskeletal dynamics and junctional stability. For example, β1-integrin, which serves as a receptor for laminin α5, signals not only through RhoA but also through Rac1 and Rap1 (53, 57), both of which have vital functions in maintaining EC junction stability (58, 59). Furthermore, integrin adhesion results in the activation of FAK and Src which phosphorylate junctional VE-Cadherin and its associated catenins, leading to junctional uncoupling from the actin cytoskeleton and VE-cadherin internalisation (43, 60–63). Thus, variable endothelial mechanosignalling and cytoskeletal dynamics downstream of heterogeneous BM deposition may create localised sites where junctions undergo destabilisation in response to endothelial stimulation.

It is remarkable to note here that pre-determined sites of leakage are preserved in the diaphragm and trachea. Compared to intravital visualisation of leakage in response to histamine, repetitive sites of leakage detected using microspheres was lower (28% vs. 14%), but still significantly different compared to random leakage site selection. This disparity may result from microsphere movement during the 24 hour interval period or differential EC response between luminal versus abluminal stimulation. Indeed, studies have shown that apical/basal polarity can influence EC response and signalling in response to VEGFA, however whether histamine signalling may similarly differ is unknown (64). Compared to the dermis, the trachea has a much higher sensitivity to histamine stimulation, highlighting the organotypic differences between these tissues. Similar to the dermis although, the trachea displayed a high proportion of pre-determined leakage sites. However, statistical analysis of the exact prevalence of pre-determined sites in the trachea is precluded by the fact that Cldn5 expression and its correlation with leakage permissiveness in this organ is currently unknown. Both the dermis and trachea appeared to retain extravasated microspheres at, or at least near sites of leakage as observed by the similar number of leakage sites following 1^st^ and 2^nd^ injections (Figures 4Bii and 5Aii). The diaphragm however showed an often significant reduction in 1st versus 2nd leakage sites, either through EC sensitisation following initial histamine stimulation, or through microsphere movement over the 24 hour interval due to distinct BM composition or mechanical pressures. Regardless, of those initial leakage sites that could still be identified, repetitive leakage sites constituted a large proportion, 50-75%. Thus, pre-determined sites of leakage highlight a functional intra-vessel heterogeneity that is common to multiple vascular beds.

In conclusion, here we demonstrate that between multiple organs, intra-vessel heterogeneity exists that define pre-determined sites of leakage where barrier breakdown occurs preferentially and to a greater degree in response to stimulation. Such sites occur due to subtle differences in the underlying BM with decreased laminin α5 deposition impacting on the junctional localisation of VE-Cadherin. Important questions remain however as to how heterogeneous deposition of BM components is established. To what degree heterogeneous EC transcriptomes are responsible for this and is patterning developmentally established or dynamically responsive to changing environments? The intra-vessel heterogeneous nature of ECs is becoming more and more realised. For example, individual ECs within vessels have been found to possess a more mesenchymal, stem-cell like phenotype and to allow the formation of vascular malformations through clonal expansion (65–69). Whether such ECs may also mark sites of pre-determined leakage is unknown. However, the emergence of highly sensitive single-cell technologies may allow such questions to be answered and holds promise for the characterisation of such intravessel heterogeneity through higher resolution mapping of the vasculature (13, 70).

## Acknowledgements

We thank Lydia Sorokin (University of Muenster, Germany) for providing the anti-Rat laminin α5 antibody.

This study was supported by the Swedish Research Council (2015-02375), Swedish Cancer foundation (CAN2016/578), Knut and Alice Wallenberg foundation project grant KAW 20150030, Wallenberg Scholar grant (2015.0275), Swedish Society for Medical Research (SSMF; 201912), an EMBO Long-term fellowship (ALTF 923-2016) as well as a Fondation Leducq transatlantic network of excellence grant in neurovascular disease (17 CVD 03).

## Author Contributions

Conceptualization, M.R and L.C.W; Methodology, M.R; Investigation, M.R and L.V; Writing, M.R and L.C.W; Funding Acquisition; M.R and L.C.W; Resources, L.C.W

## Materials and Methods

### Animals

Wild-type C57BL/6J mice age 8–18 weeks were used, unless otherwise indicated. *In vivo* animal experiments were carried out in accordance with the ethical permit provided by the Committee on the Ethics of Animal Experiments of the University of Uppsala (permit 6789/18). Each experiment was conducted on tissue from at least three animals on at least three different experimental days representing individual biological repeats. Sample size (number of acquired movies and number of mice) were chosen to ensure reproducibility and allow stringent statistical analysis. Mice were anaesthetised by intraperitoneal injection of Ketamine-Xylazine (120 mg/kg ketamine, 10 mg/kg Xylazine) to a surgical level. For intravital imaging experiments, mice were maintained at a body temperature of 37°C. For experiments using a 24 hour interval, mice regained consciousness between injections.

### Growth factors, reagents and antibodies

Rat anti-CD31 (BD Biosciences, 553370), goat anti-VE-Cadherin (R&D systems, AF1002), goat anti-collagen IV (Millipore, AB769), rat anti-laminin α5 (Sorokin et al. 1997 (71)) isolectin-B4 647 (Molecular probes, 132450), donkey anti-goat 488 and 555 (Invitrogen, A-11055 and A-21432) and donkey anti-rat 488 and 647 (Invitrogen, A-21208 and Jackson immunoresearch, 712-606-150) were used for immunostaining. For *in vivo* EC stimulation recombinant VEGFA164 (100-20; PeproTech), concentration 100 ng/μl, and histamine (Sigma-Aldrich, H7125) concentration 10 ng/μl, were used. For live imaging studies of the mouse ear dermis, a volume of approximately 0.1 μl VEGFA or histamine was injected. For vascular leakage visualisation, 2000 kDa FITC-Dextran (Sigma-Aldrich, FD2000S) or fluoromax fluorescent particles (ThermoFisher; B200, G200, R200) were used. For ligand tracing, 10 kDa TRITC-dextran was used (ThermoFisher, D1817).

### Intravital imaging and leakage site categorisation

Intravital imaging of the mouse ear dermis with intradermal injection has been described previously (3). Briefly, following systemic administration of 2000 kDa FITC Dextran mice were anaesthetised and the ear secured. Time-lapse imaging was performed using single-photon microscopy (Zeiss LSM 710) and a high N.A water-immersion lens (CF175 apochromat 25xW N.A1.1, Nikon) and used to visualise intradermal injection, traced using 10 kDa TRITC Dextran, and subsequent vascular responses.

Leakage sites were identified in time-lapse imaging, following agonist injection, as defined sites of concentrated dextran in the extravascular space. Leakage sites in response to VEGFA and histamine at the doses used occurred approximately 1 site/100 μm. Repetitive leakage sites were defined as occurring less than 5 μm from a previous site while unique leakage sites were more than 5 μm.

### Microsphere leakage assay, leakage site categorisation and BM quantification

Mixtures of histamine with fluorescent microspheres (75 μl; 1% solids) were injected systemically 24 hours apart. Five minutes following the second injection mice were perfused before dissection, immunostaining and analysis.

Leakage site categorisation as green, red or yellow was performed using a Matlab script following fluorescence analysis along vessels lengths as shown in Figure 4A. Variations in vessel widths were accounted for using the line width tool in imageJ (average vessel lengths analysed: diaphragm, 20 mm; ear, 43 mm; trachea, 76 mm). Yellow leakage sites were assigned to regions where red and green microspheres were localised within 5 μm of each other. Mean fluorescent intensities of collagen IV and laminin α5 at these sites and sites that demonstrated no leakage within the same vessel segment were normalised to isolectin B4 before comparison between each other.

### Analysis of laminin α5:VE-Cadherin correlation

Laminin α5 and VE-Cadherin fluorescence intensity was quantified along vessel lengths as shown in Figure 4A. Variations in vessel widths were accounted for using the line width tool in imageJ. To allow for variability in the number of junctional elements at any one cross-sectional site, VE-Cadherin values were normalised to the number of VE-Cadherin positive pixels within the cross section. To allow for expression differences, comparison was only made within, and not between, vessel segments. Vessel segments typically ranged between 100 – 250 μm in length.

### Immunofluorescent staining

Tissues were fixed through perfusion with 1% Paraformaldehyde (PFA) followed by immersion in 1% PFA for 2 hours at room temperature. Tissues were blocked overnight at 4°C in Tris-buffered saline (TBS), 5% (w/v) BSA, 0.2% Triton X-100. Samples were incubated overnight with primary antibody in blocking reagent, followed by washing in TBS, 0.2% Triton X-100 and incubation with appropriate secondary antibody for 2h at room temperature in blocking buffer before washing and mounting in fluorescent mounting medium (DAKO). Images were acquired using a Leica SP8 confocal microscope.

### Statistical analysis

Data are expressed as mean ± S.D. The principal statistical test used was the Students’ *t*-test. *p*-values given are from independent samples analysed by two-tailed paired or unpaired *t*-tests where appropriate. Rate of leakage was compared using linear regression and ANCOVA. Correlations were calculated using a Spearman’s Rank correlation before normalisation of multiple r values using Fisher’s Z transformation. All statistical analyses were conducted using GraphPad Prism. A *p*-value <0.05 was considered statistically significant and significances indicated as *p* < 0.05 (*), *p* < 0.01 (**), and *p* < 0.001 (***).

## Supplemental Information

**Figure S1. Repetitive vessel leakage in response to VEGFA. Relating to Figure 1.** (Left) Representative image of vessel segment susceptibility to repetitive VEGFA injection. Red and green highlighted vessels show vessel segments that leak following the first or second injection of VEGFA respectively, yellow highlighted vessels show vessel segments that leak following both injections. (Right) Quantification of vessel segment leakage (repetitive (yellow), unique 1st (red)) as a percentage of those seen following the first injection of VEGFA, with a 1 hour interval. n ≥ 6 mice, ≥ 2 acquisitions/mouse. Error bars represent mean and SD.

**Figure S2. Quantification of leakage in response to VEGFA and histamine. Relating to Figure 2.** Graph showing leakage sites per 100 μm in response to VEGFA (100 ng/μl) and histamine (10 ng/μl). Error bars represent mean and SD.

**Figure S3. Leakage sensitivities in response to repetitive VEGFA and/or histamine injection. Relating to Figures 1 and 2**. **A-D.** Graphs showing mean distances to leakage sites from the site of injection, left, and distribution of all leakage sites from the site of injection, right, following repeated VEGFA injection, 1 hour interval (A), histamine injections, 1 hour interval (B), VEGFA then histamine injection; 1 hour interval (C) or histamine then VEGFA injection; 1 hour interval (D). n ≥ 4 mice with ≥ 2 aquistions/mouse.

**E** Bar chart showing the ratios between the number of leakage sites following the first (red) and second (green) injections of histamine with a 1 or 24 hour interval (p = 0.463), alternatively, VEGFA followed by histamine or histamine followed by VEGFA, each with a 1 hour interval. n ≥ 4 mice with ≥ 2 acquisitions/mouse.

Error bars represent mean and SD. Statistical analysis was carried out using a twotailed unpaired or paired Students’ t-test where appropriate.

**Figure S4. Leakage properties at unique and repetitive sites. Relating to Figure 3. A.** Graphs showing lag periods of repetitive and unique leakage sites following the first and second injections of VEGFA with a 1 hour interval, VEGFA 24 hour interval, histamine 1 hour interval, histamine 24 hour interval, VEGFA-histamine and histamine-VEGFA, both latter cases with a 1 hour interval. n ≥ 3 mice with ≥ 2 sites/mouse.

**B.** Graphs showing degree of leakage of repetitive and unique leakage sites following the second injection of VEGFA with a 1 hour delay, VEGFA 24 hours delay, histamine 1 hour delay, histamine 24 hours delay, VEGFA-histamine and histamine-VEGFA, both latter cases with a 1 hour delay. Lines of best fit for the slopes between leakage initiation and leakage termination are shown as black lines. n ≥ 4 mice with ≥ 2 acquisitionsImouse.

Error bars represent mean and SD. Statistical analysis was carried out using an F-test for equality of variance (A) or linear regression and ANCOVA (B).

**Figure S5. Laminin α5 expression in the mouse ear dermis. Relating to Figure 4.** Representative image of laminin α5 in the microvasculature of the mouse ear dermis. Progression of the vascular hierarchy from arterioles to post-capillary venules is shown by the dashed line and arrows. Note the decrease in laminin α5 expression as arterioles branch into post-arterial capillaries and capillaries before increasing in post-capillary venules where deposition is patchy.

**Figure S6. Tracheal leakage in response to histamine. Relating to Figure 5.** Representative image of 200 nm microsphere leakage in response to 12.5 mg/kg histamine stimulation.

**Movie 1. Leakage in response to VEGFA** Time series showing the intradermal injection of VEGFA (red) in the mouse ear dermis and resulting induction of leakage from distinct sites as visualised by extravasation of circulating 2000 kDa FITC Dextran (green). Data corresponds to Figure 1B

**Movie 2. Repetitive leakage in response to VEGFA with a 1 hour interval** Time series showing leakage in the mouse ear dermis from the same microvascular region in response to repetitive intradermal injection of VEGFA with a 1 hour interval. Leakage is visualised through extravasation of 2000 kDa FITC Dextran (pseudocolour). Data corresponds to Figure 1C.

**Movie 3. Leakage profiles of repetitive and unique leakage sites in response to VEGFA** Time series showing leakage in the mouse ear dermal microvasculature following the intradermal injection of VEGFA. Leakage is visualised through extravasation of 2000 kDa FITC Dextran (pseudocolour). Repetitive and unique sites were identified through a subsequent injection of VEGFA. Data corresponds to Figure 3B.

**Movie 4. Leakage profiles of repetitive and unique leakage sites in response to histamine** Time series showing leakage in the mouse ear dermal microvasculature following the intradermal injection of histamine. Leakage is visualised through extravasation of 2000 kDa FITC Dextran (pseudocolour). Repetitive and unique sites were identified through a subsequent injection of histamine. Data corresponds to Figure 3C.

**Movie 5. Leakage in response to tail-vein histamine with a 4 hour interval** Time series showing leakage in the mouse ear dermal microvasculature following repetitive tail-vein injection of histamine as visualised through extravasation of 2000 kDa FITC Dextran (pseudocolour). The first injection (left) and second injection (right) took place with a 4 hour interval. Note that no leakage takes place following the second injection. Histamine injections took place at 20 seconds in both movies.

**Movie 6. Leakage in response to tail-vein histamine with a 24 hour interval** Time series showing leakage in the mouse ear dermal microvasculature following repetitive tail-vein injection of histamine as visualised through extravasation of 2000 kDa FITC Dextran (pseudocolour). The first injection (left) and second injection (right) took place with a 24 hour interval. Note that leakage takes place following the second injection and often from the same sites. Histamine injections took place at 20 seconds in both movies.

